# People report having consistent idiosyncratic ‘diets’ of imagined sensations when they re-experience the past, and pre-experience the future

**DOI:** 10.64898/2025.12.12.694050

**Authors:** Derek H. Arnold, Loren N. Bouyer, Blake W. Saurels, D. Samuel Schwarzkopf

## Abstract

To some extent, humans can re-experience the sensations of past events and pre-experience the future. These capacities are inter-related. But there are substantial individual differences. At the extremes, small minorities of people report that they either cannot have imagined experiences at all, or that their imagined sensations are as real to them as their actual experiences of the physical world. We wanted to know if such individual differences are uniform across different types of imagined experience (e.g. vision, audio, taste and smell), or if people generally have idiosyncratic patterns of different types (vision, audio, taste and smell) of imagined experiences. We find that people report having idiosyncratic ‘diets’ of different types of imagined sensation, characterised by differences in salience. One person might have more salient imagined visual than taste experiences, while another reports the reverse. Moreover, these propensities are consistent across people’s attempts to re-experience the past, and to pre-experience the future, and they predict people’s experience and usage of different types of imagined sensation in their everyday lives.

---

People have a capacity to re-experience the sensations of past events, and to imagine pre-experiencing the future [1]. When asked to recollect a childhood party, adults might be able to conjure a recollected sensation of hearing people sing happy birthday, they might be able to have an imagined re-experience of tasting a cake, and visualise the setting. These people could have similar imagined sensations when they plan for a future event.

Evidence suggests that people’s capacities to re-experience the past and to imagine pre-experience future events are inter-related [2]. These cognitive operations entrain activity in similar regions of the human brain [3–5], and the ability of people to recall the details of specific past events is associated with their ability to imagine the features of future events [6–7]. Episodic memories and future thoughts can also entrain affective responses of equivalent emotional intensity [8]. Conceptual similarities between episodic memories of the past, and of people’s capacity to imagine future events, has led to the proposal that a common set of cognitive mechanisms underlies both capacities. Hypothetically, this allows people to mentally travel in either direction in time, backwards into their past or forwards into an anticipated future – the mental time travel hypothesis [9].

There are large individual differences in the capacity of different people to have volitional imagined sensations, of the past or the future [10]. If we consider the capacity to visualise, for instance, evidence suggests a continuum ranging from people who cannot visualise at all [11–12], to people who feel that their visualisations are as real to them as actually seeing [13]. The term Aphantasia was initially coined to refer to people who cannot visualise [11–12], but these issues are multisensory [14]. People who report being unable to visualise will also often report being unable to have other types of imagined sensation [15–17]. For instance, some people describe themselves as being unable to experience imagined sounds [16–17], and this can extend to people who report that they cannot experience subvocal speech [18–20]. People might also report having an inability to have a range of other imagined sensations, such as of taste or of smell [14,17].

Not only do some people report having an inability to have different types of imagined sensation [14, 16–17], there is evidence that these different types of imaginative inability can cluster. A recent study considered the experiences of a large number of people who reported having an inability to visualise. Of these, ∼30% reported having no other type of imaginative inability (pure visual aphantasia), and ∼24% reported being unable to have any type of imagined sensation (multi-sensory aphantasia). The authors were also able to identify groups of people who reported that they could only have imagined sensations in a single sensory modality (e.g. they might only have imagined audio sensations), and who reported having an inability to have imagined sensations in just two sensory modalities [e.g. visual and kinaesthetic, see 16].

Visual Aphantasics generate fewer episodic details than controls when they contemplate both past and future events [10], and they tend to report having idiosyncratic patterns of inability to conjure different types of imagined sensation [16]. It therefore seems reasonable to hypothesise that people who report being unable to have imagined sensations will tend to report having a common idiosyncratic diet of imagined sensations when they re-experience the past, and pre-experience the future – although this has never been formally tested.

We wanted to know if such tendencies would be echoed in the general population – who have in common a capacity to have different types of imagined sensation (e.g. of vision, audio, taste and smell, [see 17, 21]). Will people who are able to have different types of imagined sensation nonetheless have idiosyncratic experiences of these different sensations, perhaps characterised by differences in relative salience? Moreover, we wanted to see if any such propensity would shape people’s experience and usage of different types of imagined sensation in their daily lives. If a given type of imagined sensation is more salient to you than other types of imagined experience, will you more likely have that type of imagined sensation in your daily life? There is good evidence that this type of relationship holds for visual imagery, with people who report typically having vivid imagery also reporting that they often visualise in their daily lives [22]. However, we do not know if people generally have idiosyncratic experiences of different types of imagined sensation (e.g. vision, audio, taste and smell), or if the strengths of these imagined experiences will shape the likelihood of people having and using these different types of imagined experience in their daily lives. Moreover, if the mental time travel hypothesis is valid, and people rely on a common set of cognitive mechanisms to recall the past and to imagine the future [2, 9], any imaginative idiosyncrasies should be consistent across people’s attempts to re-experience the past, and to pre-experience the future.

## Results

### What proportion of our sample were Aphantasic?

Of our participants, 1% obtained the lowest possible vision subscale score of 0 (see Figure 1) on the Plymouth Sensory Imagery Questionnaire (PSIQ) [23]. This instrument provides a subjective assessment of the salience of people’s imagined sensations, with vision, audio, taste, smell, touch and body movement subscales. In addition to vision, 1.3% of our participants had the lowest possible scores on the smell and body movement subscales, and just 0.4% had the lowest possible scores on the audio, taste, and touch subscales.

**Figure 1.**
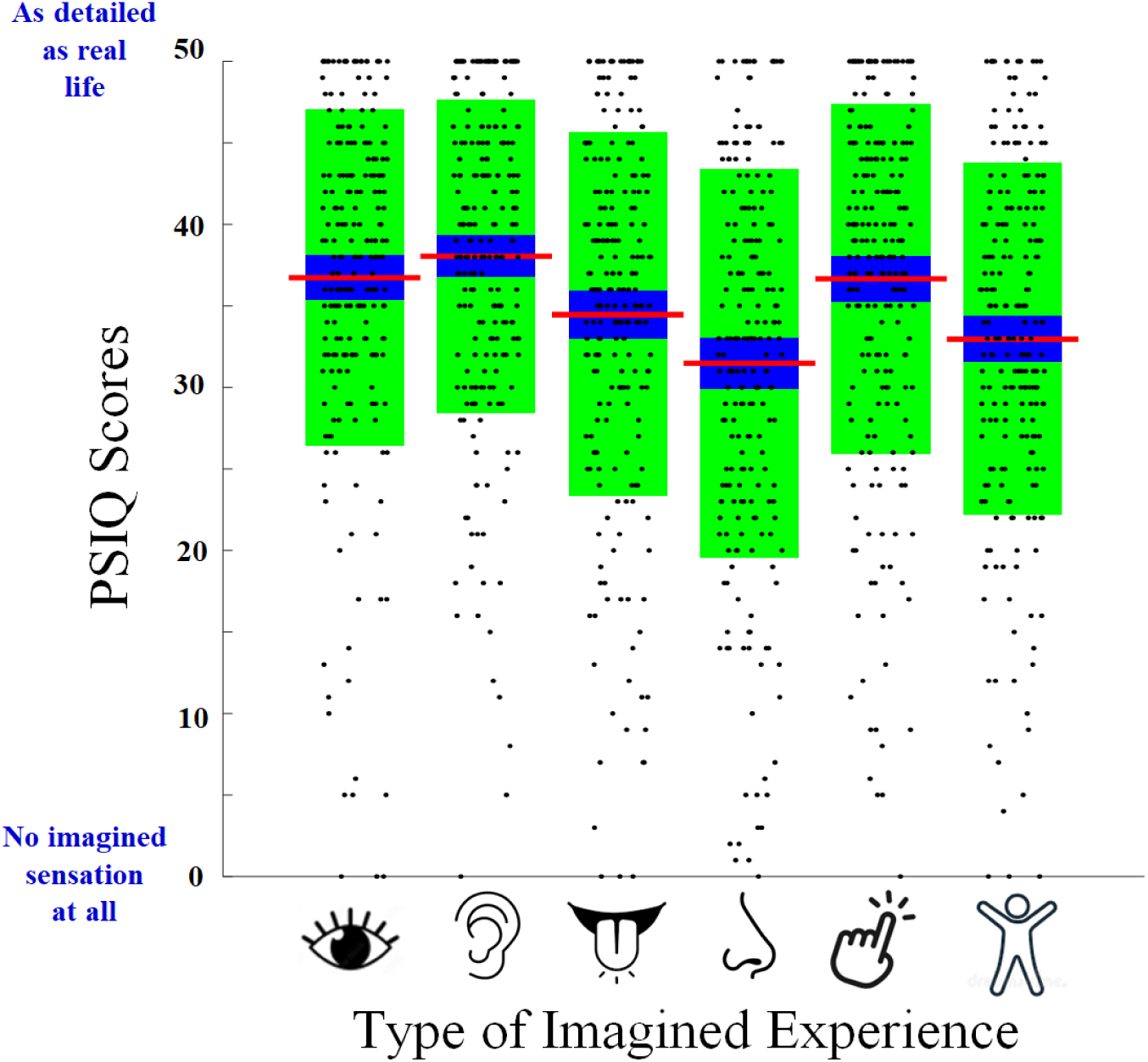
Boxplots of PSIQ subscale scores. Red horizontal bars depict average scores. Blue regions depict +/- 1 SEM from the average, and green regions depict +/- 1 SD from the average. Black dots are individual data points, with a random X-axis jitter. Subscale scores of 0 indicate that the participant reported that they had no imagined sensation at all across all subscale items, and scores of 50 indicate that the participant reported that their imagined experiences had seemed as detailed as real life across all subscale items. Icons below the horizontal axis represent the different subscales (visual, audio, taste, smell, touch and body sensation).

We regard minimal possible scores as evidence of Aphantasia – an inability to voluntarily have different types of imagined sensation [11–12]. These estimates are commensurate with other estimates of the prevalence of visual aphantasia in the general population (∼0.7% [24–25]). This means that the overwhelming majority of people in our sample reported having a capacity to have all the different types of imagined sensation we have investigated, scoring moderately across PSIQ subscales.

### Prevalence of people who did not report having different types of imagined sensation when thinking about specific future or past events

While our data suggest that most people in our sample *could* have all the different types of imagined sensation we have targeted for investigation (visual, audio, taste and smell), this does not dictate that they *would* have all of these types of imagined sensation each time they are prompted to think about novel future and specific past events [see 14]. In Figure 2 we have plotted the proportions of people who reported that they did not have a particular type of imagined sensation when they were cued to think about specific past or potential future events. These data highlight that an overwhelmingly majority of our participants reported having all of the types of imagined sensation we have investigated. Just 4% or less of people reported that they had *not* had a particular type of imagined sensation when prompted. The only exception was that ∼11% of people reported that they did not have an imagined re-experience of past smells (see Figure 2). So, not only did our participants report that they *could* have the different types of imagined sensation we have investigated, the vast majority reported that they *did* have all of these types of imagined sensation – both when they were asked to think about specific events from their past and when they were asked to think about potential future events.

**Figure 2.**
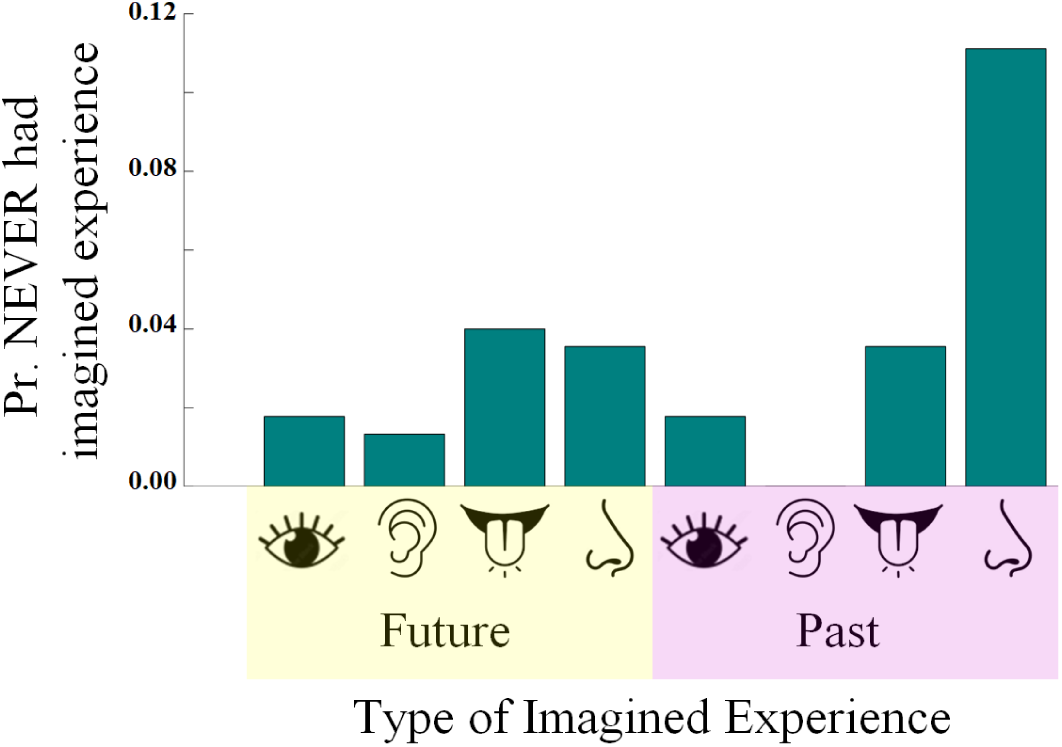
Bar plot of the proportions of people who reported not having different types of imagined sensation, when either imagining a novel future event (left side, yellow shaded icons) or when recollecting a specific past event (right side, pink shaded icons).

### People report having imagined sensations that have idiosyncratic relative strengths, that are consistent across thoughts about the future and the past

A key question we wanted to address is whether people would report having idiosyncratic patterns of imagined experiences, that are consistent across attempts to re-experience the past and to pre-experience the future.

We conducted sequential linear regression analyses – to predict people’s average strength ranking scores for each type of imagined sensation (vision, audio, taste and smell) when they thought about specific past events. In each case predictors in the first model were average rank order scores for all the types of imagined sensation we had measured, minus those for the same type of imagined experience when people had thought about novel future events (so there were 6 predictors in the first model, and 7 in the final). For example, visual rank order scores for future events would be left out as predictors of visual rank order scores for past events in the first model, and added as the 7th set of predictors to a second and final model. In 3 of 4 cases the final model explained >5% of additional variance in people’s average imagined strength rank scores relative to the first model (8% to 34%; see <u>Supplemental Table1</u> for a depiction of these data). The only exception was the final model for past audio events. These data reveal special relationships in-between imagined sensation strength rank scores for the same types of imagined sensation, that relate to specific past events and to potential future events. So, people who felt that smell had been the strongest of their imagined sensations when they were thinking about a potential future event were likely to report that this had been their strongest imagined experience when they thought about a specific past event. As a direct test of this proposition, we additionally conducted a non-parametric shuffle test procedure [e.g. 26-28].

For this analysis, we adjusted average rank order scores into individual rank orders, separately for data relating to novel future and to specific past events. If two or more modalities had an equal average top ranked score, they were each given a ranking of #1. The next highest scored modality (or modalities) would then be ranked 3, and so on. If two or more modalities had average rank order scores of 5, these would be ranked equal last – 4.

We identified what numbers of people had given each type of imagined experience the same ranking, regardless of whether it had been triggered by thinking about a novel future or by a specific past event (see Figure 3). We then conducted 500 iterations, wherein we randomly interchanged past rankings in-between different participants, and determined how many pairs of unshuffled future and shuffled past rankings were then matched (i.e. are both 1, or are both 3) for the same type of imagined sensation (vision, audio, taste or touch). Doing this 500 times creates a distribution of the number of times that matched rankings would happen at chance in the dataset, if there had been no correspondence between future and past rankings at an individual level. Note that this treatment of data accounts for generalized trends, as more matches would occur between unshuffled and shuffled datasets if particular rankings were common. An overall P-value is calculated from how many of the 500 sets of chance pairings had contained the same or a greater number of matched rankings relative to the original unshuffled dataset.

**Figure 3.**
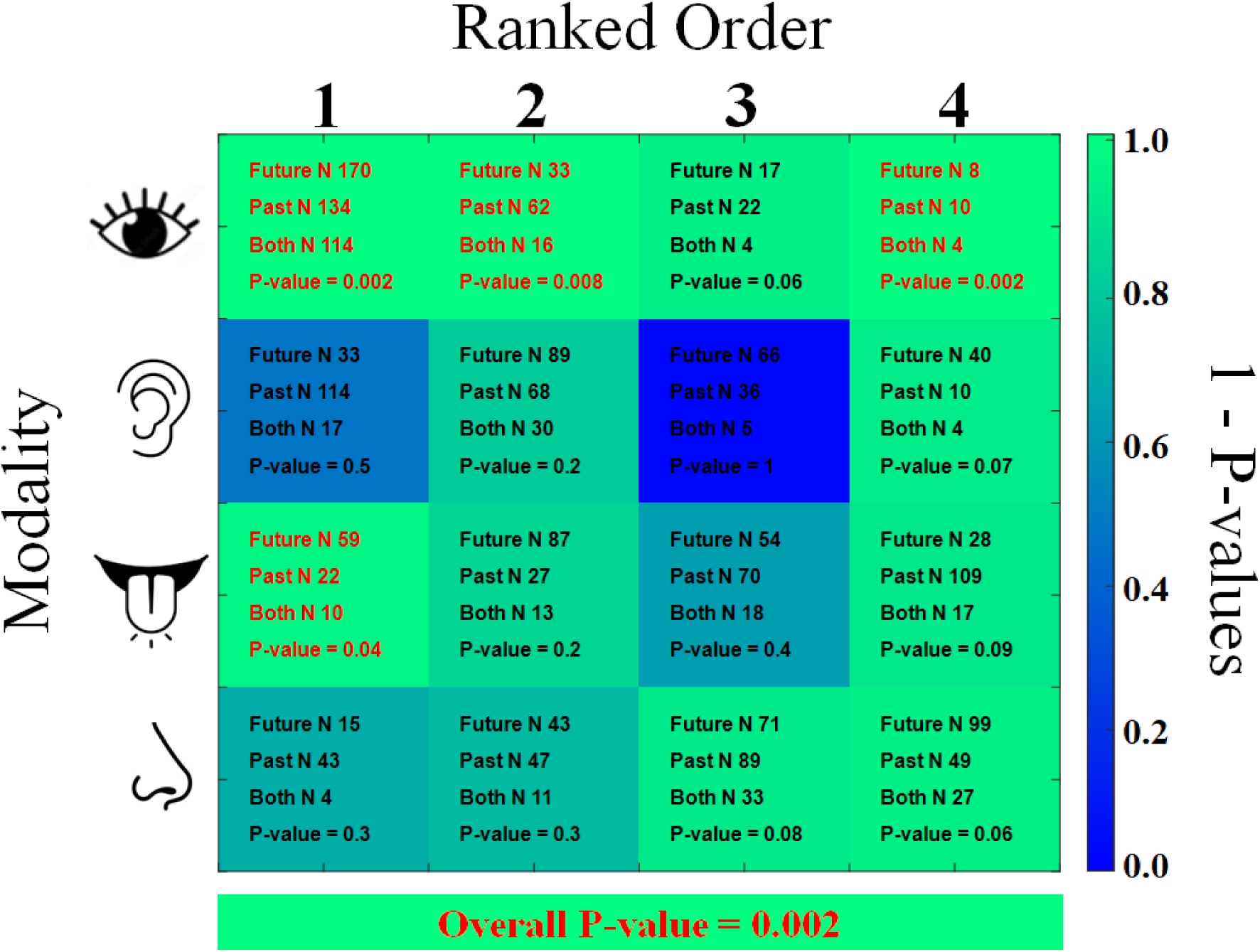
Heatmap describing 1 - P-values. The different types of imagined modality are arranged along the Y-axis. The rank orders that people could give to describe the strength of each type of imagined experience are arranged along the X-axis. Within each cell, the number of participants who gave the relevant type of imagined experience that ranking when thinking about future events is shown above, the number of participants who gave that ranking when thinking about past events is shown in the middle, and the number of participants who gave that ranking *both* when thinking about future *and* past events is shown below. Colour coded 1-P-values indicate what proportion of 500 ‘*shuffles*’ of past rankings resulted in the same or in more matches in-between unshuffled future and shuffled past rankings relative to the original, unshuffled dataset. See main text for further details.

To prevent p-values of 0, the actual number of matched rankings is added to the shuffle test chance distribution, so the minimum possible p-value that can result from this procedure is 0.002 – and this was the overall P-value that was delivered by this testing procedure (see Figure 3, bottom). This reveals that, overall across all 4 tested modalities for this analysis, people’s rankings of the relative strengths of their imagined sensations were both idiosyncratic but consistent across their thoughts about specific past and potential future events.

Our shuffle test procedure delivers an overall statistic, indicative of how likely it is that we could have obtained the total number of matched future / past rank orders, for the same types of imagined experience, if there had been no association between future and past order rankings at an individual level. To illustrate which combinations of imagined sensation (vision, audio, taste or smell) and order rankings (1-4) had driven this overall association, we repeated this type of analysis separately for each of the 16 potential combinations of imagined modality and rank orders. These analyses revealed that subsets of people reliably gave the same rankings to visual events (see Figure 3, top row), and a subset of people reliably ranked taste as their strongest type of imagined sensation (see Figure 4, 3^rd^ row from top).

**Figure 4.**
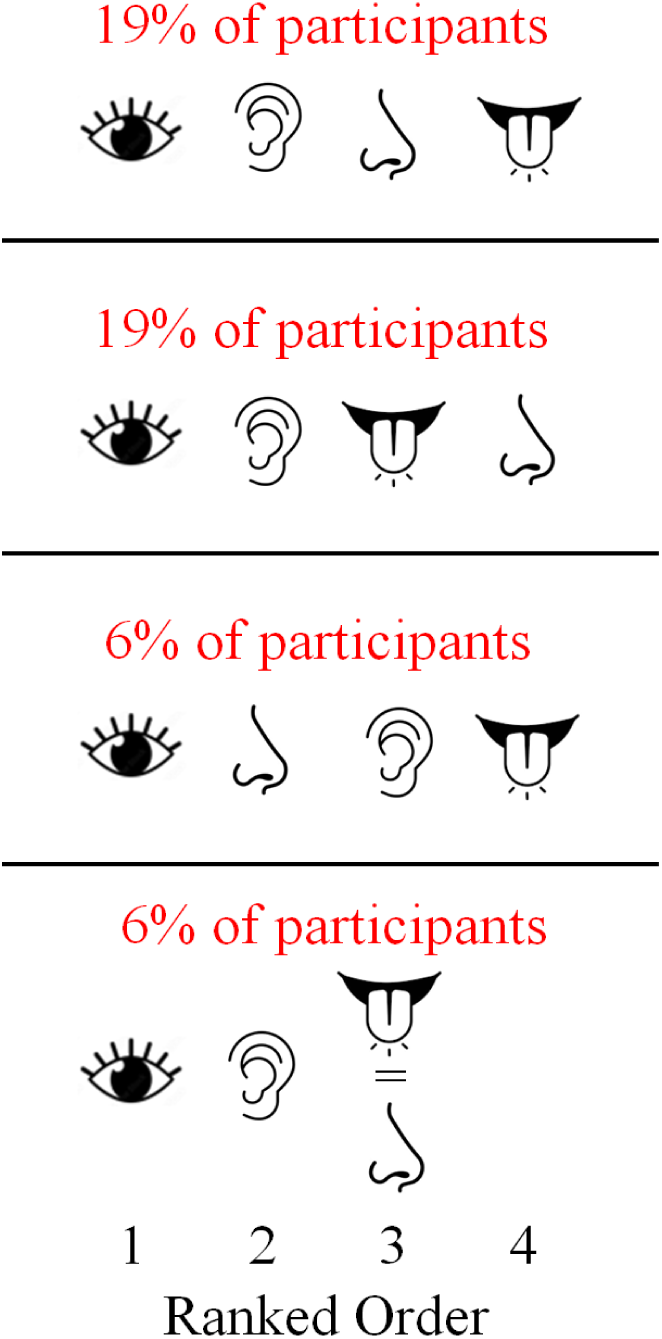
Graphics depicting the 4 most common sets of rank orders, in terms of the strengths of different types of imagined experience, for data combined across imagined experiences elicited by thinking about novel future and specific past events.

So far our analyses have established the existence of imaginative idiosyncrasies. In addition to these, however, our data speak to some rank order combinations being more common than others across people. For this analysis we first averaged people’s rank order scores across future and past events, to create a set of combined rank order scores for each participant. We then converted these into a set of combined rank orders for each individual (i.e. from strongest – 1, to weakest – 4). We were then able to calculate how many different combinations of combined rank orders people had reported. There were 48 in total (a total number that is possible because combinations of the 4 sensory modalities could be equally ranked). This provides further evidence of idiosyncratic rank ordering. However, some combinations were far more common than others.

In Figure 4 we have depicted the 4 most common combinations of combined rank orders. Across all four, vision was ranked the strongest type of imagined sensation. Of these 4 combinations, the top two were each reported by ∼19% of our participants, and each of these had audition ranked as second. These top two sets of combined rank orders differed only in terms of whether taste or smell had been ranked as the weakest type of imagined sensation. The third most common set of combined rank orders (∼6% of participants) had smell ranked second, audition third and taste last. The fourth most common (∼6% of participants) had audition ranked second, and taste and smell equal third. These top 4 combinations of combined rank orders were reported by ∼50% of our participants (see Figure 4). We have reported on these common combinations to highlight that there are similarities across people, in addition to the imaginative idiosyncrasies we have detected.

### Analyses of people’s general impressions of the frequency of different types of imagined experience about the future and the past

In addition to having people report on the content of their thoughts when they were cued to think about specific novel future and past events, we asked them to report on their general impressions of how often they have different types of imagined sensation when thinking about the future and the past (see Supplemental Figure 2).

We conducted sequential linear regression analyses on these data to predict people’s frequency scores for each type of imagined sensation about the past. Similar to our last set of sequential linear regression analyses, in each case predictors in the first model were average frequency scores for each type of imagined sensation we had measured, minus scores for the same type of imagined experience when people thought about future events (for example, vision scores for future events were omitted as predictors of vision scores for past events, and so on for all 8 types of imagined experience). In the majority of cases final models explained >5% of additional variance in people’s imagined frequency scores (8% – 23%; see <u>Supplemental Table2</u>). The exceptions were final models for Taste, Smell and Touch.

### Do people’s imaginative idiosyncrasies impact their daily lives?

We have established that people report having imaginative idiosyncrasies, characterised by differences in the relative salience of different types of imagined sensation – that tend to be consistent across people’s attempts to re-experience the past, and to pre-experience the future. We now turn to the question of whether these imaginative idiosyncrasies impact people’s daily lives.

Our bespoke SISS questionnaire provides 4 subscale scores (see Figure 5) that are indicative of how often people experience and use different types of imagined sensation (vision, audio, taste and smell) in their daily lives. Subscale questions include items asking people to report on how often they visualise places they plan to visit, rehearse conversations in their mind before speaking, have an imagined sensation of the taste of potential shopping items, and imagine how they will smell before they exercise (amongst other items – for further details see Supplemental material, Spontaneous Imagined Sensations).

**Figure 5.**
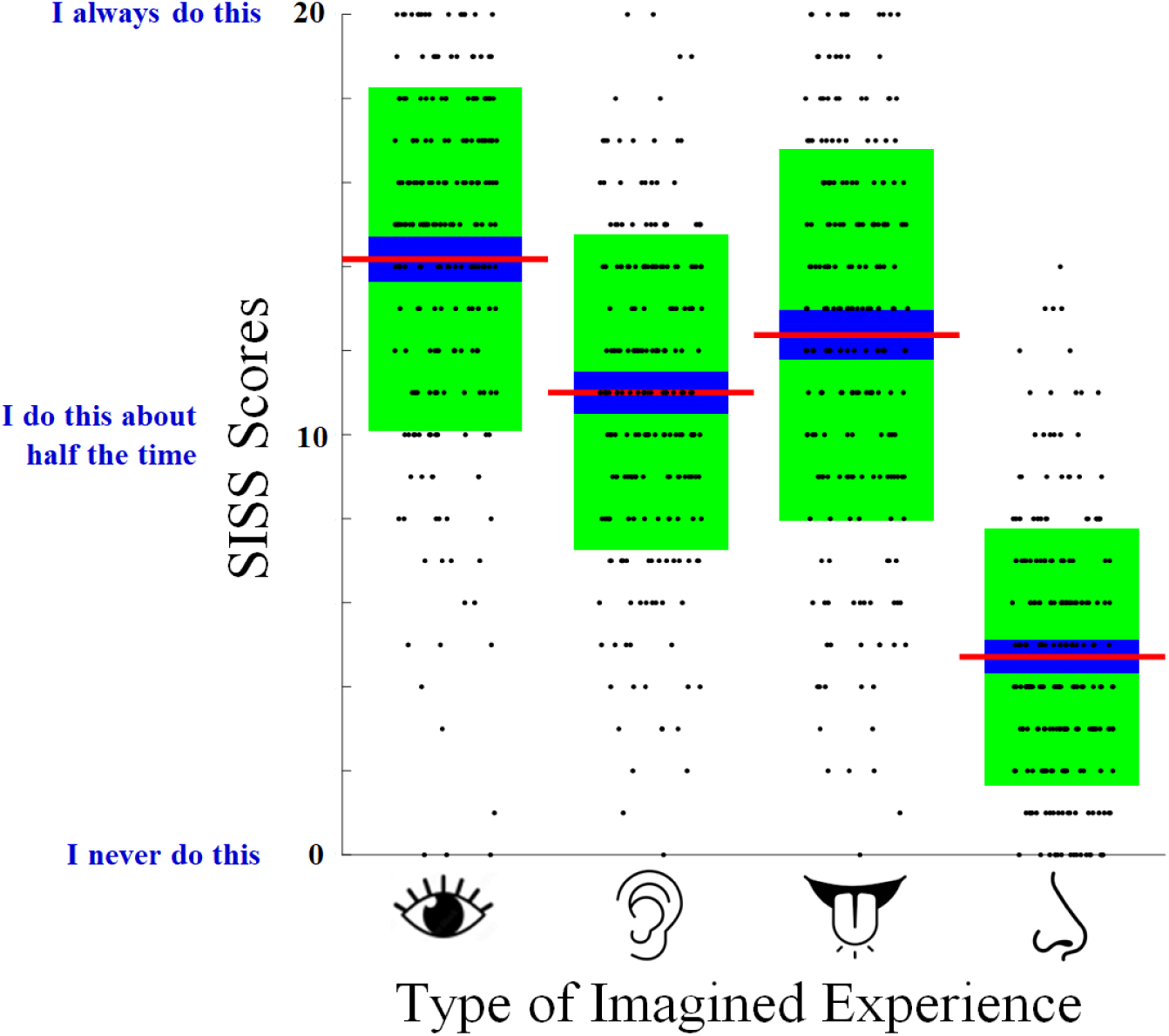
Boxplots of people’s SISS subscale scores. Details of the plots are as described for Figure 1.

Three of the four SISS subscales (Vision, Taste and Smell) are useful psychometric instruments. These have good levels of internal consistency, with respective Cronbach’s alphas of 0.76, 0.82 and 0.73. Moreover, our results capture some evidence for the convergent validity of these SISS subscales, as each was positively associated with the corresponding PSIQ subscale score (spearman’s Rho’s of 0.37, 0.47 and 0.29 respectively, p-values < 0.00625; see Figure 5, left column). Note that in all correlational analyses that we report on, outlier datapoints (> +/- 3 S.D.s from the conditional mean) are excluded from analyses, and we conduct Shapiro-Wilk tests to assess if contributing datasets are both consistent with a normal distribution. This was never true of any of our datasets, so we calculate Spearman’s rank correlation coefficients, which we report on as ‘rho’ in figures and in text. In this set of analyses, we tested for 8 relationships, so the nominal significance of associations are evaluated against a Bonferroni adjusted alpha level of 0.00625.

The PSIQ is a self-report instruments that measures the subjective salience of people’s typical experiences of different types of imagined sensation – inclusive of imagined sensations of vision, taste and smell [23]. The positive association of SISS and PSIQ subscale scores therefore provides evidence of convergent validity, as it is reasonable to assume that the salience of different types of imagined sensation (as given by PSIQ subscale scores) might be related to the likelihood of these types of imagined sensation being experienced and used by people in daily life [23] – as given by SISS subscale scores.

Positive associations between SISS and PSIQ subscale scores suggests that people are more likely to report experiencing and making use of a particular type of imagined sensation when it is salient to them. SISS subscale scores (Vision, Taste and Smell) were also associated with people’s average rank order scores (with respective spearman’s Rho’s of -0.19, -0.22 and - 0.25, p-values < 0.00625; see Figure 6, right column). These relationships are *negative* because low rank order scores indicate that type of imagined sensation was rated as *stronger* relative to other types of imagined sensation (see Supplemental Figure 1). These data therefore reveal that people were more likely to report experiencing and making use of a particular type of imagined sensation when they felt it had been more salient to them than other types of imagined sensation.

**Figure 6.**
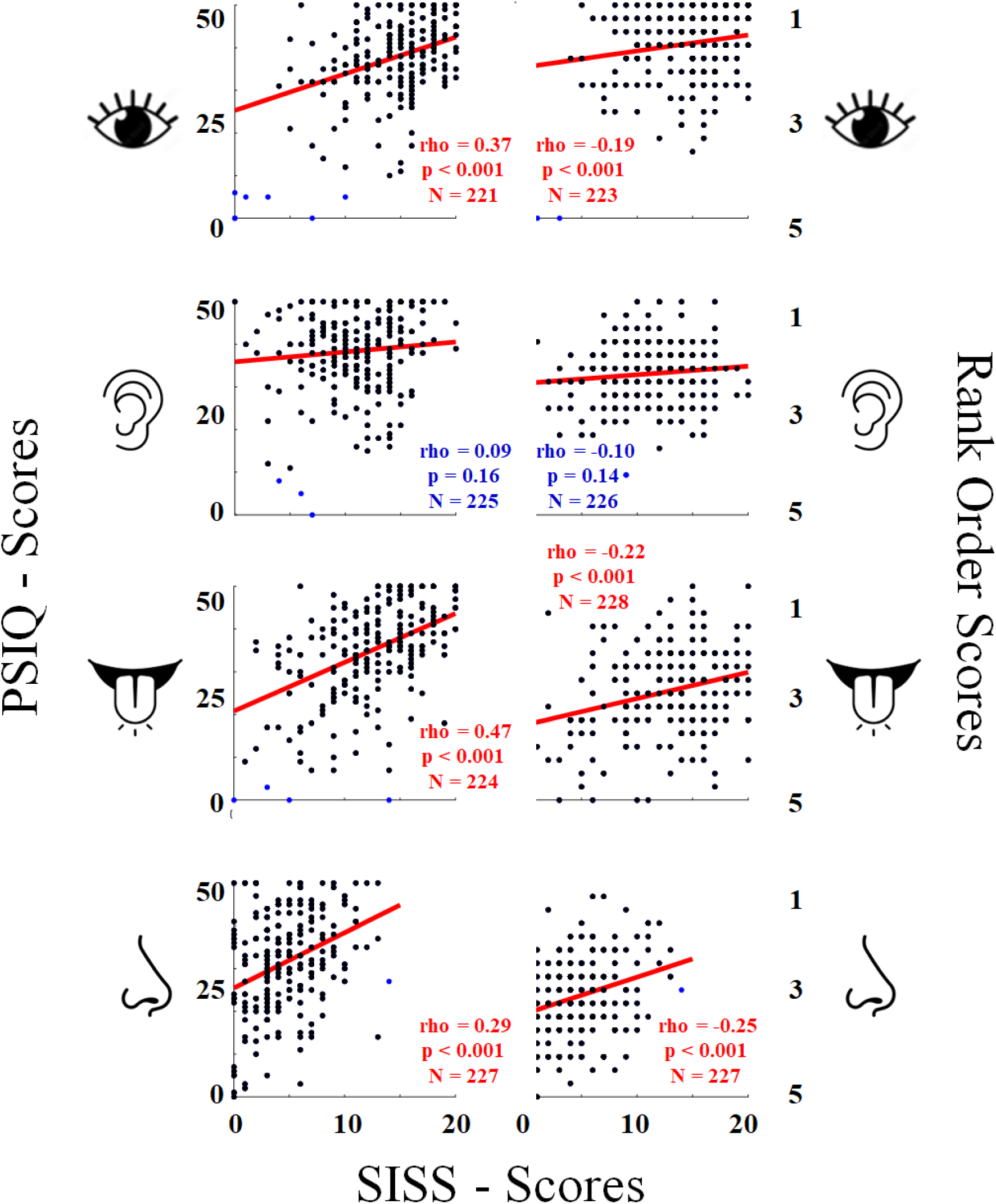
X/Y scatterplots of people’s SISS subscale scores (X-axes) and PSIQ subscale scores (left column, Y-axes) and average Rank Order Scores (right column, Y-axes). Note that Y-scale labels are reversed for average Rank Order Scores, as low scores indicate that these types of imagined sensation were ranked stronger than other types of imagined sensations.

## Discussion

Our data show that people report having an idiosyncratic diet of different types of imagined sensation, characterised by differences in relative salience (see Figure 3 and Supplemental Figure 1). These idiosyncrasies were consistent across people’s attempts to re-experience past events, and to pre-experience novel future events (see Figure 3 and Supplemental Table 1). Moreover, people’s imaginative idiosyncrasies predicted how often they would report experiencing and using different types of imagined sensation in their daily lives (see Figure 6 and Supplemental Table 2). Our data therefore suggest that people generally have imaginative idiosyncrasies, characterised by differences in the salience of different types of imagined sensation, and these shape their experiences of daily life.

### People report having idiosyncratic ‘diets’ of imagined sensations

Previously, there had been good evidence for individual differences in the salience of particular types of imagined experience [23] – especially for visual imagery [12–15]. There was also evidence that the vividness of a person’s visual imagery was related to the probability that they would report visualising in daily life [22]. We have expanded on these findings, to show that people report having idiosyncratic ‘diets’ of different types of imagined sensation, characterised by distinctive variance in salience across different types of imagined sensation (e.g. Vision, Audio, Taste and Smell), and that these imaginative idiosyncrasies shape people’s experiences of daily life.

An important feature of our data is that it suggests that imaginative idiosyncrasies are not just a matter of academic interest. Rather, these seem to have an impact on people’s daily lives. For example, people were more likely to report that they visualise places they plan to visit if they had reported that their imagery is typically vivid – and if they rated their visual imagery as more salient to them than other types of imagined experience. There were similar contingencies in relation to the probabilities that people would report that they imagine what different items will taste like when shopping, and that they imagine how they might smell when they think about exercising (see Figure 6).

### Imaginative idiosyncrasies in the general population and Aphantasia

Our data are the first to suggest that people generally have idiosyncratic ‘diets’ of different types of imagined sensation – characterised by differences in relative salience (see Supplemental Figure 1 and Figure 3). This echoes Aphantasia research. Studies have shown that there are identifiable sub-groups of Aphantasics [16]. By definition, all Aphantasics are unable to visualise – but a proportion of Aphantasics report that they cannot have *any* imagined experiences, whereas other subgroups report having different combinations of imaginative inabilities [16].

Almost all the people in our study reported that they could have each of the different types of imagined experience we have investigated (see Figure 1). But people’s reports also revealed that the relative strengths of the different types of imagined experience they could have were both idiosyncratic (see Figure 3) and were reliable across attempts to re-experience the past, and to pre-experience the future (see Supplemental Tables 1-2). So, imaginative idiosyncrasies do not just relate to a uniform modulation of imagined salience. Rather, these can relate to distinctive patterns of difference in the relative salience of the various types of imagined sensations that people can have. Moreover, these differences do not just manifest at the extremes of an imaginative spectrum, but are rather evident across the population.

#### Some differences in the salience of imagined sensations are common across people

Across samples of the general population, visualising has consistently been estimated to be more common and more salient to people than are other types of imagined experience [14, 22–21, 28–32]. Our data are consistent with the pre-eminence of visual imagery, in terms of the strength of imagined sensations (see Figures 3, 6 and 7). Just 28% of people rated another type of imagined experience as being stronger than vision. It was also common in our data for people to rate audio as a relatively strong imagined sensation, and to rate taste and smell as weaker imagined experiences (see Figure 1 and Supplemental Figure 1). Our data highlight idiosyncrasies in the precise ordering of these different types of imagined sensation (see Figure 5), but they also capture some common general trends.

#### The SISS audio subscale needs to be revised

Our data suggest that 3 of the 4 SISS subscales are potentially useful psychometric instruments. The Vision, Taste and Smell subscales had good levels of internal consistency (each having a Cronbach’s alpha > 0.7) and have evidence of convergent validity, as each was positively associated with the corresponding subscale of another validated instrument – the PSIQ [23] (which measures the salience of different types of imagined sensation, see Figure 1). The audio SISS subscale was a notable exception. It had poor internal consistency (Cronbach’s alpha 0.57) and was unrelated to PSIQ audio subscale scores (see Figure 6).

The lack of internal consistency across SISS audio subscale items could be due to these measuring different psychological constructs. People’s inner voices might be distinct from their capacity to have other types of imagined audio sensation, as there are reports of people who have an inner voice, but cannot have other types of imagined audio experience [19–20]. So, for example, questions about imagined rehearsals of conversations might not be strongly associated with questions about imagining hearing the voices of family members. There is also an issue that SISS audio questions could measure involuntary imagined experiences (i.e. when I hear or read certain names or phrases, it makes me imagine hearing a song). We have previously found that the incidence of involuntary audio experiences of cued scenarios (i.e. do not think about the sound of a toilet flushing) are not predicted by the clarity of people’s typical experiences of imagined audio sensations [36]. Whatever the cause, the SISS audio subscale is not currently a useful psychometric instrument, and it needs to be revised.

#### There did not have to be an association between the salience of people’s imagined sensations and their experiences of daily life

Our data suggest that the salience of different types of imagined sensation, and the strength of these relative to other types of imagined experience, predict if people will report having and using that type of imagined experience in their daily lives (see Figure 6). To speculate about cause, it is equally plausible that people who have more salient imagined sensations will more often choose to have and use these experiences, or that the frequent experience and use of a given type of imagined experience will overtime reinforce underlying processes and so strengthen that type of imagined experience. Readers might feel that this type of relationship is intuitive, but from a conceptual perspective this type of relationship did have to eventuate.

Almost all people can see, but there are vast differences in the clarity of people’s vision. People do not *see* less often if their visual acuity is poor. Similarly, people could have imagined experiences of diverse strength, but have these equally as often, or at idiosyncratic frequencies that are unrelated to the strengths of their imagined sensations. Our data suggest a different relationship that might be more intuitive, but this relationship was not inevitable.

#### Our data relate to people’s subjective experiences

Our data unambiguously relate to the *subjective* experiences of people’s thoughts, and to how these relate to people’s reported experiences of daily life. Our results show that people report having idiosyncratic patterns of different types of imagined experience, characterised by differences in salience, that are consistent across their thoughts about the future and the past. Moreover, these differences are related to the types of imagined experience people report having and using in their daily lives. The pertinent question then becomes ‘*why do people report having imaginative idiosyncrasies, that seem to impact their experiences of life*? People typically have good insight into their subjective experiences [46–48], so the most straightforward interpretation of our data would be that people have different experiences of their imagined sensations, and they accurately report on these. This would explain why these measures seem to be related to everyday experiences. It remains possible, however, that people’s imagined experiences are all similar, and that some other explanation must be found for the reliable differences in people’s reporting on their imagined sensations. Future studies can assess these possibilities by examining if self-reported individual differences, in terms of the salience of different types of imagined sensation, can successfully predict tertiary measures of imagined sensations, inclusive of their neural substrates [e.g. 34, 36 – 39].

#### Why would there be an association between people’s attempts to re-experience the past, and to pre-experience the future?

The suggestion that the capacity to re-experience the past, and to pre-experience the future, are intertwined might surprise some readers. There was, however, good pre-existing evidence that encouraged this hypothesis [2–5]. For instance, these cognitive operations entrain activity in similar brain regions [3–5], and the ability of people to recall past events is associated with their ability to imagine future events [7]. We would further suggest that, for most people, attempts to pre-experience the future are likely informed by re-imagining features of past events [49–50].

#### Conclusions

Our data show that people report having idiosyncratic patterns of different types of imagined sensation, characterised by differences in relative salience, that are consistent across attempts to re-experience the past and to pre-experience the future. Moreover, these imaginative idiosyncrasies predict what types of imagined sensation people report having and using in their daily lives.

## Methods

All data relating to this study are publicly available via UQ eSpace.

### Ethics

Ethical approval was obtained from the University of Queensland’s (UQ) Ethics Committee. The experiment was performed in accordance with UQ guidelines and regulations for research involving human participants. Each participant provided informed consent to participate, and they were informed that they could withdraw from the study at any time without prejudice or penalty.

### Participants

A total of 289 people participated in this study. Of these, 61 failed one or more attention check items. These were trivially easy items, such as ‘*If you are paying attention, please select the middle option.*’ After these exclusions, our final sample consisted of 228 people, 121 of whom identified as female, 105 as male, and one each who identified as *other* and *prefer not to say*. The average age was 43 (S.D. 13) ranging from 19 to 76. All participants reported having English as their first spoken language. Participants were recruited via the Prolific platform (www.prolific.com) and were paid £4.50 to participate.

The sample size was dictated by resource availability. We had sufficient funds to pay for 300 respondents. This would have delivered ∼0.938 power to detect weak associations (r = 0.2). While 300 payments were made, only 289 responses were recorded on the Qualtrics platform, and 61 of these people failed attention check items. The final sample of 228 people should deliver ∼0.862 power to detect weak (r = 0.2) associations.

#### Stimuli and Procedure

This study was conducted online, via Qualtrics (Qualtrics, Provo, UT). After providing informed consent to take part, participants were first asked to respond to some demographic questions (age & gender identity).

After responding to demographic questions, participants completed the visual, audio, smell, taste, touch, and bodily sensation subscales of the PSIQ [22]. These provide a subjective assessment of the salience of a participant’s imagined sensations to cued scenarios (e.g. *‘imagine the appearance of a sunset’*, or *‘imagine the sound of the sound of children playing’*). Subscale scores range from a minimum of 0 (which indicates that the participant reported having ‘*No imagined sensation at all*’ in response to all 5 subscale scenarios) up to a maximum of 50 (which indicates they reported that their imagined experiences were ‘*As detailed as real life*’ in response to all 5 subscale scenarios). This questionnaire was included to establish that our participant pool was comparable to other studies of the general population, and to establish that the majority of participants felt that they could have each of the different types of imagined sensation we wanted to focus on in our investigation.

Participants then completed a bespoke questionnaire (see Supplemental Material: Experiences of Thought – Ranked Order Scale). This assessed peoples’ experiences when they were cued to think about novel future and specific past events. There were also questions about people’s general experiences when thinking about the future and the past, which quizzed them about the frequency of different types of imagined experience.

To assess people’s experiences when thinking about novel future and specific past events, people were invited to imagine events that were intended to elicit a range of different types of imagined sensation (visual, audio, taste and smell). We restricted these items to asking people about just four types of imagined experience because we had difficulty conceiving of life events that would reliably trigger more than four types of salient recallable imagined sensation. For future events, people were cued to imagine a pair of novel future events (a special dinner when they would try something new, or a trip to a beach that they had never previously been to, with someone they had never been to a beach with). For past events people were asked to imagine re-experiencing a specific event from their past (a party they had attended, or a time when they had been sick with a cough). For each of these four scenarios, participants were asked to rank order imagined visual, audio, taste and smell sensations in terms of strength, with options to rank sensations as equal, and to report that they had not experienced that type of imagined sensation at all. This provided an averaged score ranging from 1 (strongest sensation) to a maximum possible score of 5 (no sensation at all) for each combination of type of imagined sensation (visual, audio, taste and smell) and tense (*future* and the *past*).

We also asked people to think generally about their experiences when thinking about the future and the past, and to report on how frequently they have different types of imagined sensation. These included imagined sensations of hearing themselves speak, of hearing other sounds (e.g. instrumental music, bird songs or traffic noise) that they were *not* mimicking with their inner voice, of visualising, imagined tastes, smells, feelings of touch, body movement, and imagined sensations of the feel of their bodies when happy or sad. People could report that they ‘*never did this*’ (scored as 0), that they were ‘*unsure*’ (1), that they ‘*sometimes do this*’ (2), that they ‘*often do this*’ (3), or that they ‘*always do this*’ (4). This resulted in each participant having a score from 0 to 4 to describe how often they have different types of imagined sensation when thinking about the past or the future. We included more types of imagined sensation in these questions, as these types of experience did not have to be linked to a specific past or potential future event.

To assess people’s experiences and volitional use of different types of imagined sensation in their daily lives, we devised a second bespoke questionnaire – the Spontaneous Imagined Sensations Scale (SISS). Some items for the visual subscale (SISS-V) were adapted from the Spontaneous Use of Imagery Scale [22]. This last instrument was designed to address conceptually similar questions in relation to visual imagery. To our knowledge, there was no similar existing multisensory instrument, so we devised further sets of questions to quiz people about their imagined experiences of audio, tastes and smells. Across all subscales items invited people to indicate how often they have different types of imagined experience. Participants could select ‘I never do this – 1’, ‘2’, ‘I do this about half the time – 3’, ‘4’, or ‘I always do this – 5’, which were respectively scored from 0 to 4. There were 5 questions for each subscale, so SISS subscale scores can range from 0 to 20.

Full details regarding all questions asked of participants, and their responses, are publicly available from (link removed for anonymity).

## Acknowledgements

This research was supported by a Discovery Project Grant, funded by the Australian Research Council, awarded to D.H.A.

## Conflict of interest

The authors declare no competing financial interests.

## Data and materials availability

All data and analysis scripts for this project are available via [UQeSpace](https://espace.library.uq.edu.au/view/UQ:08c31c8)

## Author contributions

D.H.A conceived of the study, programmed the experiment, analysed data, created figures, and wrote the first draft of the manuscript. All other authors contributed to conceptual discussions informing the study design and edited successive drafts of the manuscript.

## SUPPLEMENTAL MATERIAL

### Average rank order scores for the strengths of imagined sensations for the past and future

In Supplemental Figure 1, we depict average rank order scores, relating to the different types of imagined sensation that people reported having when they thought about novel future and specific past events.

**Supplemental Figure 1.**
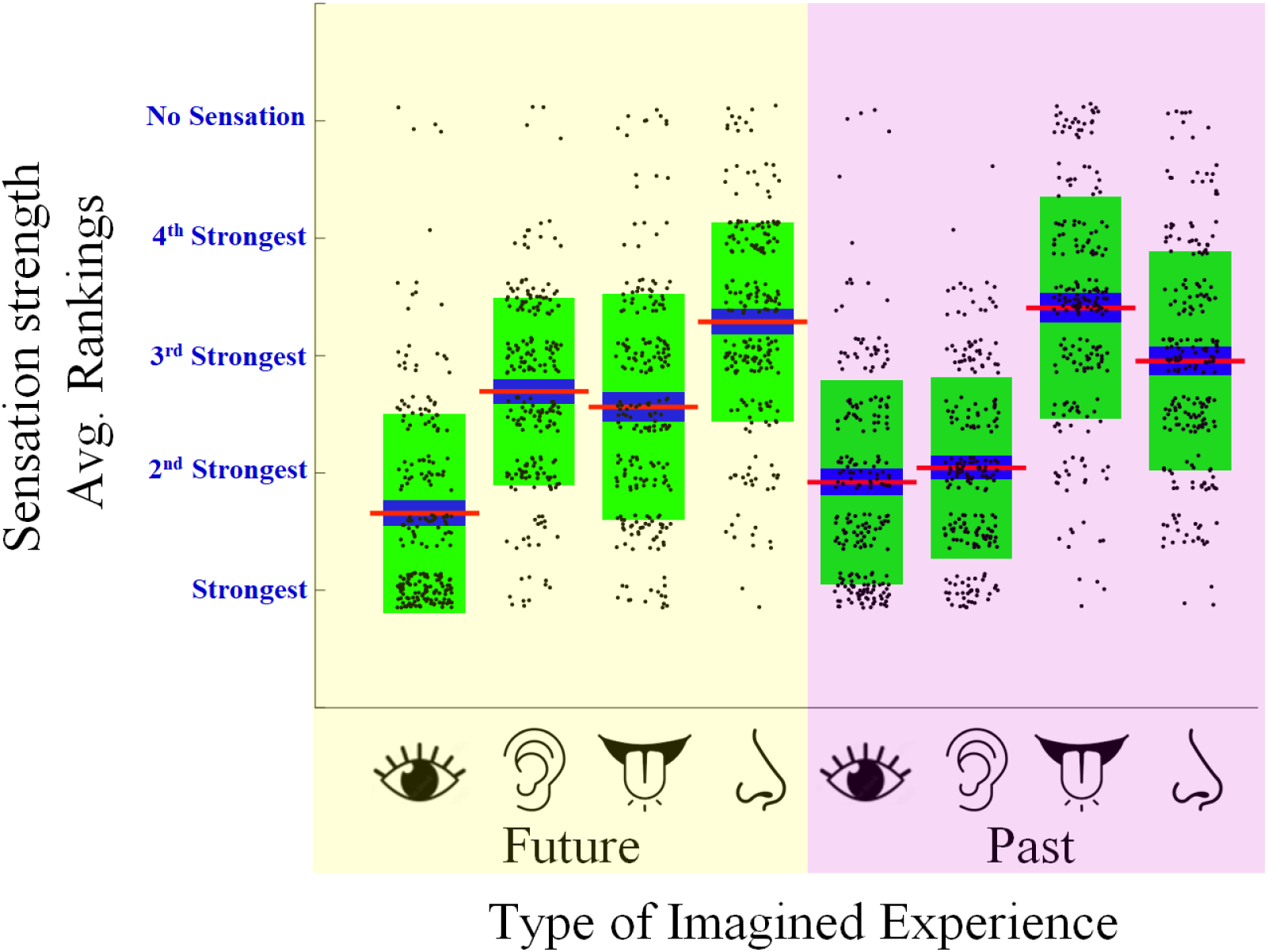
Boxplots describing people’s average rank order scores for different types of imagined sensation, experienced when people thought about specific cued future (yellow shaded data) and past (pink shaded data) events. Details of the plots are as described for Figure 1 in main text.

### Results of linear regression analyses, predicting the salience of imagined sensations for past events

In Supplemental Table 1, we depict key results of linear regression analyses, predicting the average rank order scores for different types of imagined sensation (Vision, Audio, Taste and Smell) that people had when thinking about specific past events. The beta values of predictors in the final model are depicted in columns. In each case the first model included as predictors all average rank order scores for other types of imagined sensation, minus the matching modality for future events. These were added as predictors to the second and final model.

The variance explained by each model (r^2^ values) are depicted to the right of the columns of beta values, along with the % change in variance explained (by the final model). When this exceeds 5% of additional variance explained, the change is highlighted in dark pink.

**Table 1.**
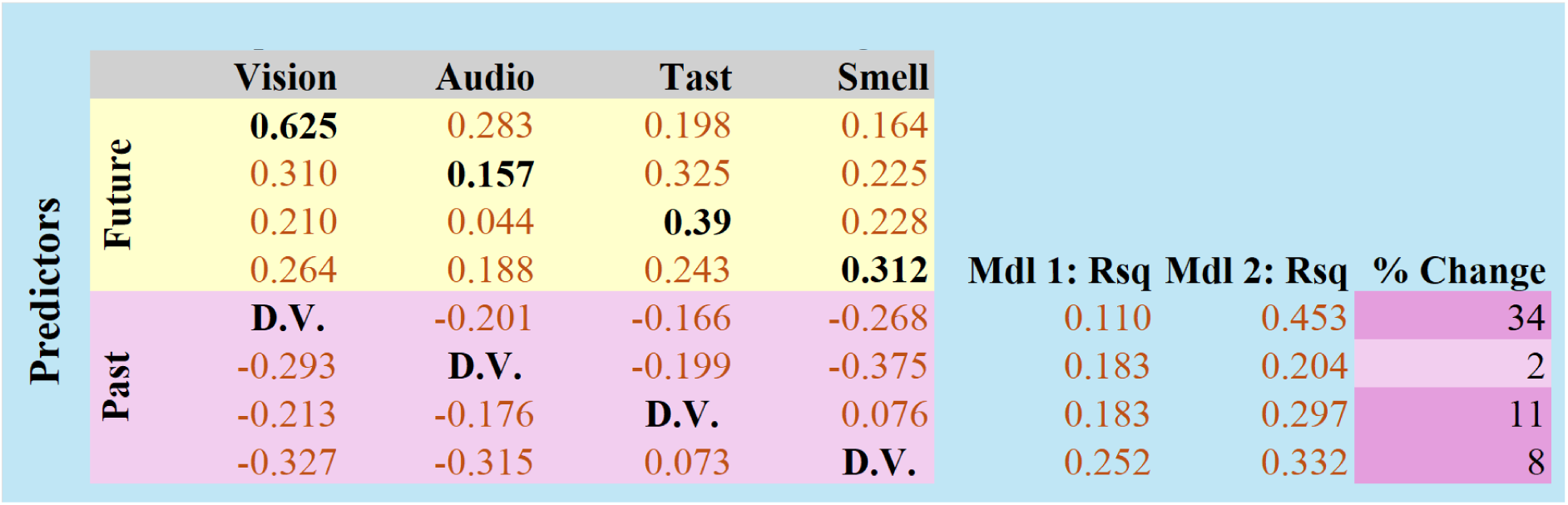
Specific Events - Final Model, Linear Regression Beta Values. Standardized Beta coefficients given to different predictors in the final model of 4 linear regression models (arranged in rows, from top). Future and Past predictors are presented separately, and are colour coded. The dependent variable for each model is signified (D.V.).

### Average frequency ratings for different types of imagined sensation for past and future events

In Supplemental Figure 2, we depict average frequency ratings, relating to the different types of imagined sensation that people reported having when they thought about novel future and specific past events.

**Supplemental Figure 2.**
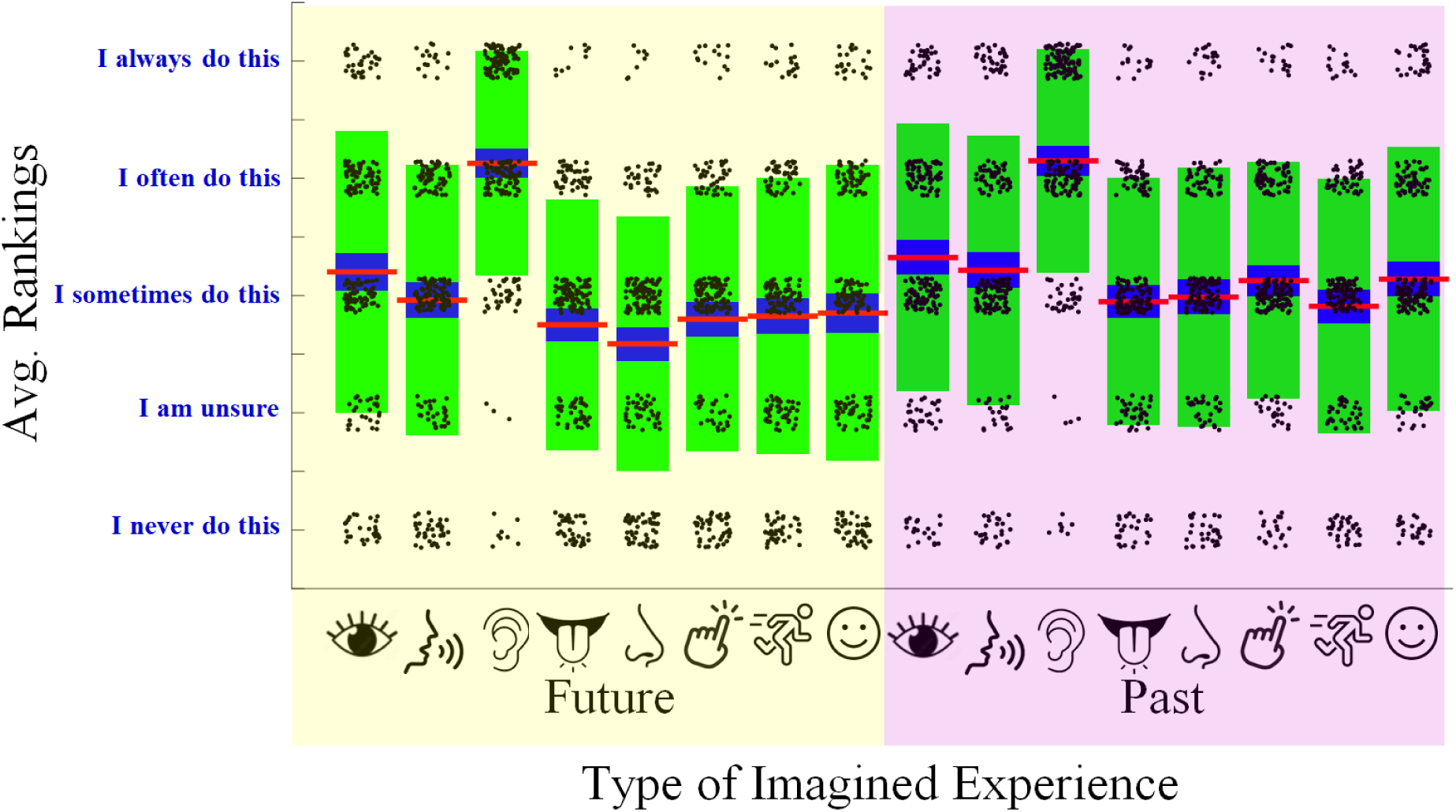
Boxplots describing distributions of people’s ratings of their general experiences when thinking about the future (yellow shaded data) and the past (pink shaded data). From left, icons depict imagined sensations of vision, inner speech, audio other than inner speech, taste, smell, touch, bodily movement and bodily sensations when feeling happy or sad. Details of the plots are as described for Figure 1 in main text.

### Results of linear regression analyses, predicting the reported frequency of different types of imagined sensation for past events

In Supplemental Table 2, we depict key results of sequential linear regression analyses predicting frequency scores for each type of imagined sensation about the past. Details of these analyses were as described for the set of linear regression analyses predicting average rank order scores.

**Supplemental Table 2.**
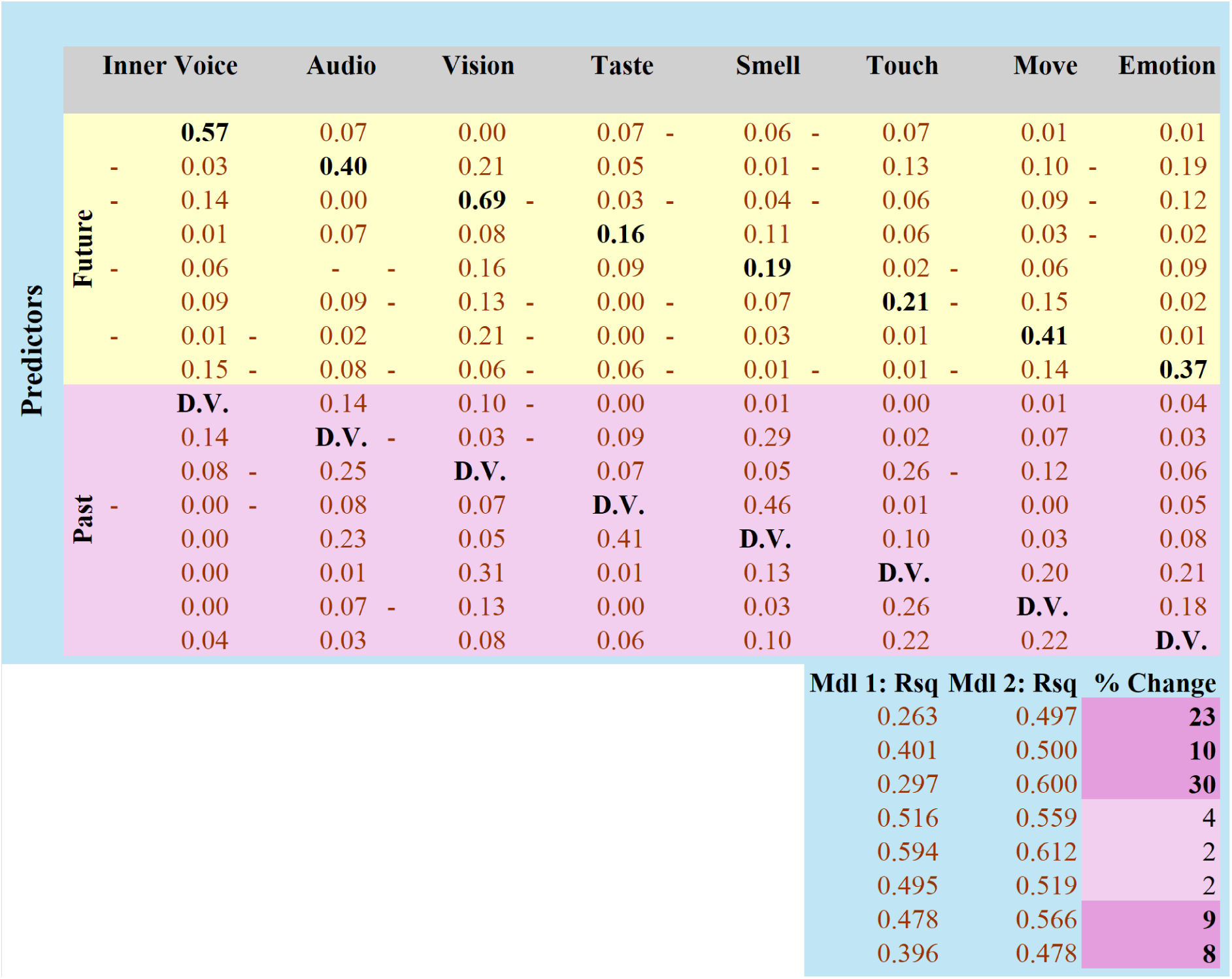
General Evants - Final Model, Linear Regression Beta Values. Beta weights given to different predictors in the final model of linear regressions, which each predicted scores indicating how frequently people reported having imagined sensations in daily life. All details are as described for Supplemental Table 1, with the exception more conditions are depicted, and model r^2^ and % change statistics are depicted below standardized Beta coefficients values

### Spontaneous Imagined Sensation Scale scores

In Supplemental Figure 3, we depict Spontaneous Imagined Sensation Scale (SISS) scores, for Vision, Audio, Taste and Smell subscales.

**Supplemental Figure 3.**
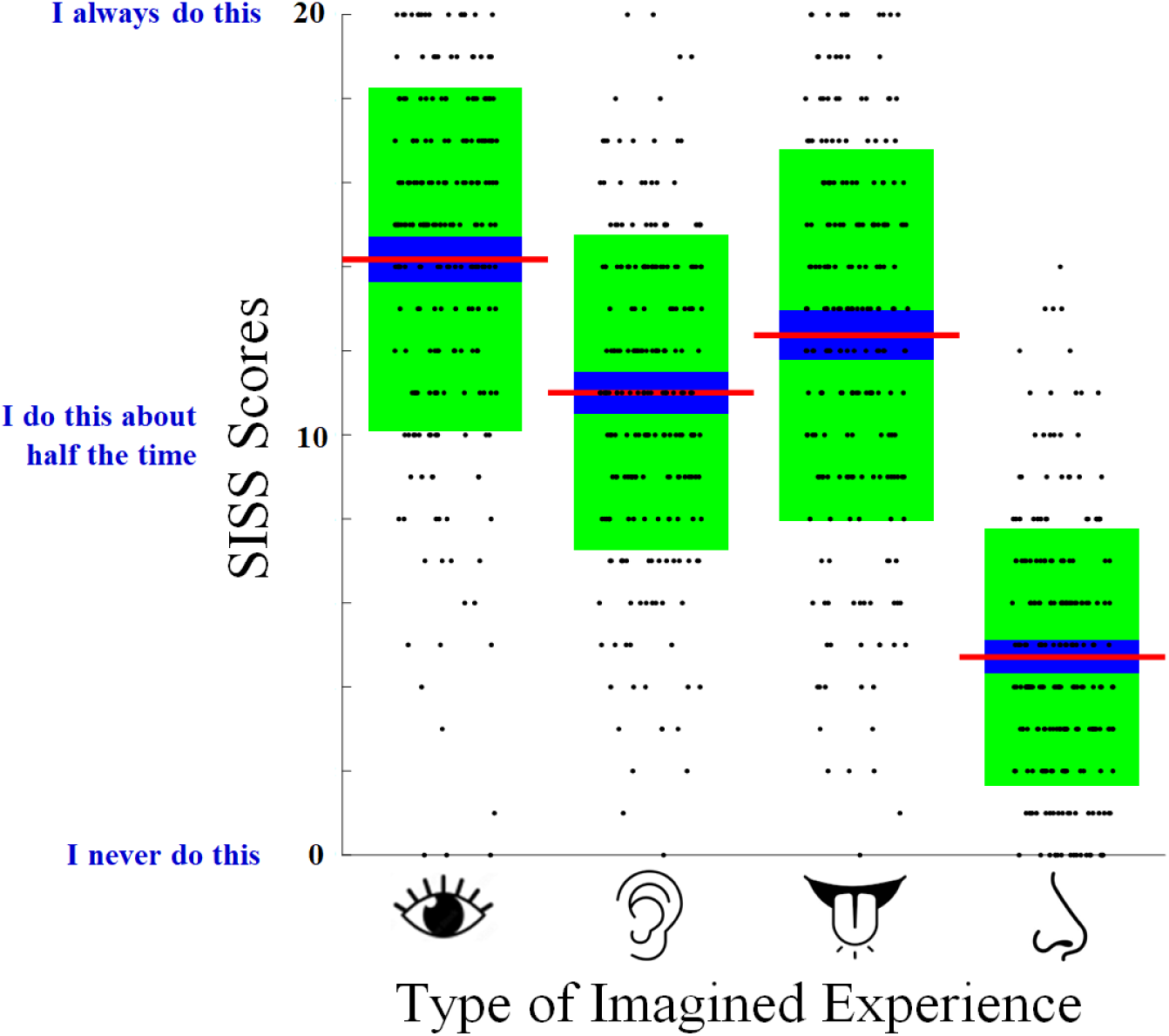
Boxplots of people’s SISS subscale scores. Most details of these plots are as described for Figure 1 in main text.

## Spontaneous Imagined Sensations Scale

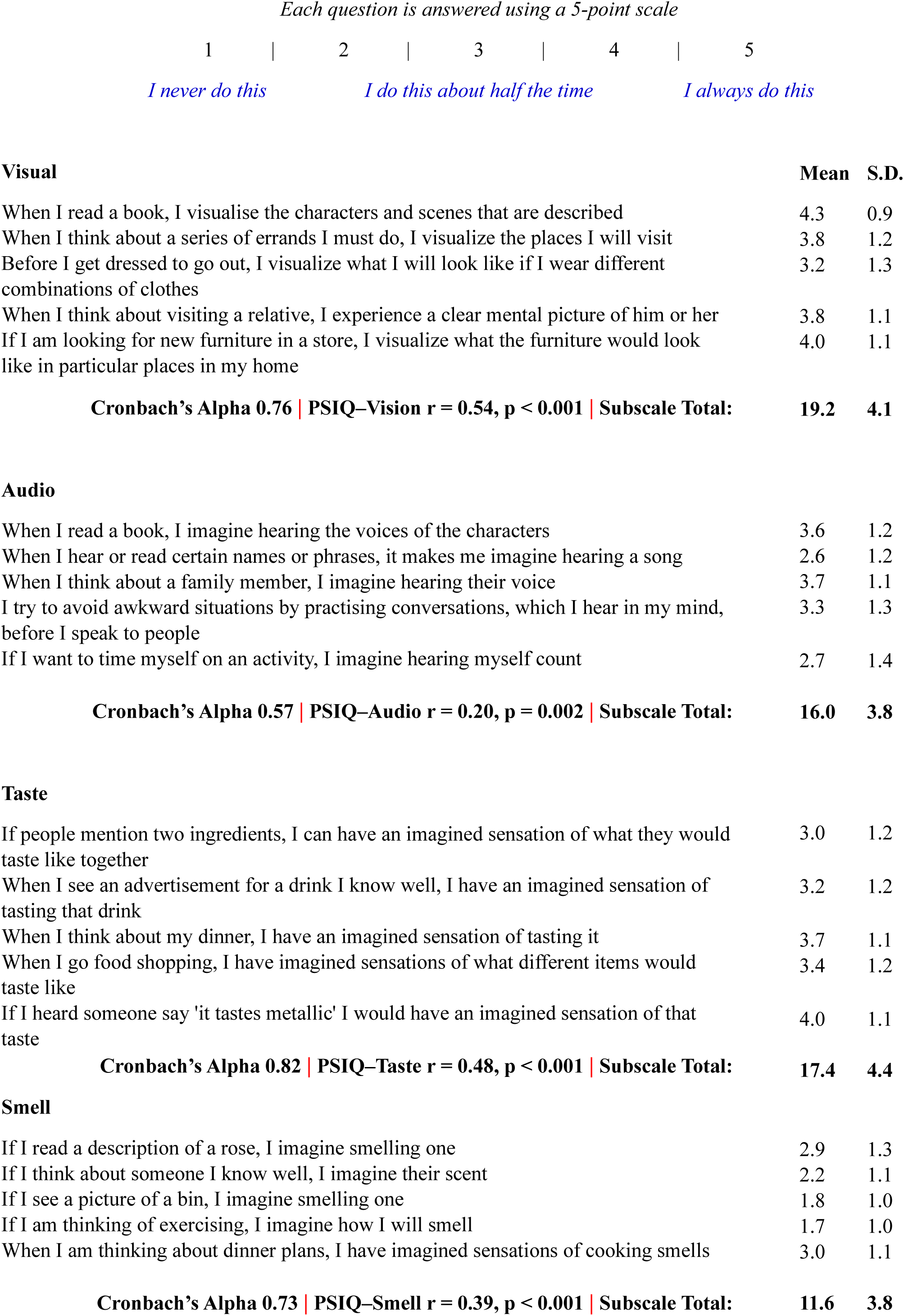

### Experiences of Thought – Ranked Order Scale

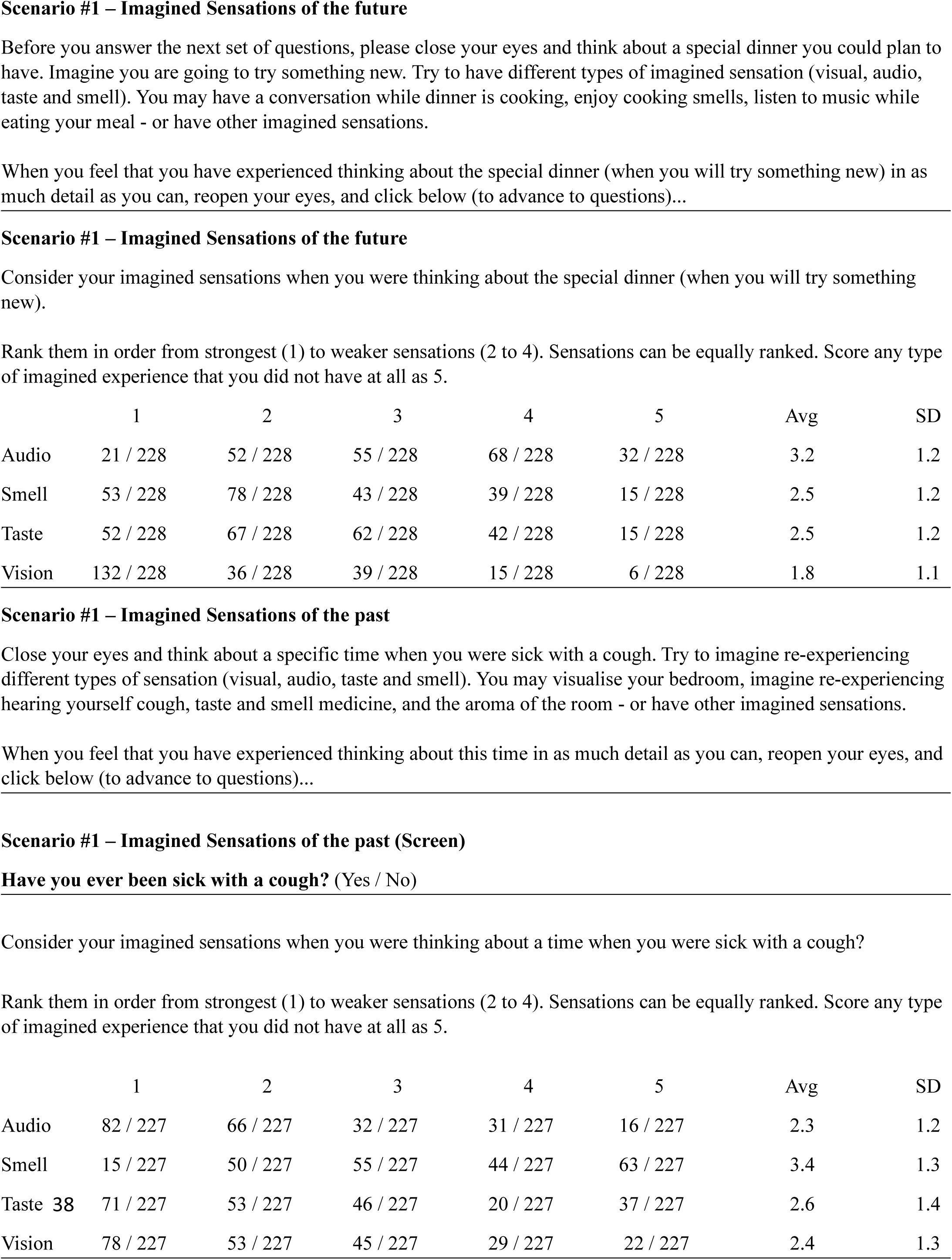

## Notes

https://espace.library.uq.edu.au/view/UQ:08c31c8

### Competing Interest Statement

The authors have declared no competing interest.

